# Interspecific trait differences drive plant community responses on serpentine soils

**DOI:** 10.1101/2024.03.08.584143

**Authors:** Guillaume Delhaye, Panayiotis G. Dimitrakopoulos, George C. Adamidis

**Affiliations:** Ecosystem Stewardship, Royal Botanic Gardens Kew, United Kingdom; Biodiversity Conservation Lab., Department of Environment, University of the Aegean, Greece; Biology Department, University of Patras, Greece

**Keywords:** Plant community, ultramafic, intraspecific variation, leaf economics spectrum, metal tolerance, environmental filtering, competition

## Abstract

1. Serpentine ecosystems are characterized by multiple environmental stressors such as high levels of trace metals such as nickel (Ni), low availability of macronutrients and low water retention. These harsh environmental conditions exert a strong selective force on the vegetation, but their effect on the functional trait composition of the communities remains unknown.
2. In 26 plots on four serpentine sites on Lesbos Island (Greece), we measured six leaf functional traits related to resource acquisition and stress resistance on the 20 most abundant plant species. We quantified the proportion of variance explained by inter- and intraspecific traits difference and tested if individual species show changes in trait values explained by soil Ni content. We investigated the selective value and the community level changes for each trait along the natural soil Ni gradient using a species multilevel model approach and functional diversity analyses. We also tested the role of the abundant serpentine endemic *Odontarrhena lesbiaca* in driving these patterns.
3. Intraspecific variation explained by soil Ni content is smaller than 2%, with most of the variance being explained by interspecific differences in trait values and most species do not show significant changes in trait values. At the community level, leaf thickness is the only trait driving an increase in species abundance along the gradient. Functional diversity analyses suggest a shift towards a stress tolerance syndrome (thick leaves with low SLA values) on Ni rich soils, but an increase in the diversity of these traits. However, these patterns are driven by the increasing abundance of *O. lesbiaca*. When this species is excluded, there is an increase in the community mean leaf area and SLA, suggesting that the community does not respond to metallic stress with classical stress syndromes.

**Synthesis.** Intraspecific variation in leaf trait responds little to soil metal toxicity. Endemic species harbour original trait values compared to species with broad distribution which should justify their conservation as a priority.

## Introduction

Species traits are related to the strategies of species living in different biological communities and demonstrate adaptation to different conditions of competition, disturbance, and stress (Donovan et al. 2011). At the organ or individual level, traits can be coordinated. For example, it was suggested that stress tolerance traits are tightly coordinated with growth traits, at both the individual plasticity and the species evolutionary levels, to produce a phenotype that is adapted to the environment (Chapin et al. 1993). Differences in soft traits can drive differences in hard traits driving specific responses to the environment (Chapin et al. 1993; Belluau and Shipley 2018). For example, the study of leaf economic traits (Wright et al. 2004) informs about the strategy deployed by plants to thrive along environmental gradients (Dong et al. 2020). The combination of species abundance, their traits and environmental conditions form the basis for studying the impact of abiotic environment on community assembly processes, such as the harsh environmental conditions produced by increased soil metal concentrations (Kazakou et al. 2008; Delhaye et al. 2016; Gervais-Bergeron et al. 2023). The three main levels at which trait variation can be studied are the intraspecific, the interspecific and the community level.

At the community level, functional diversity, i.e., the variety of ecological strategies occurring in a community, is expected to decrease in areas facing severe abiotic stress (Mason & de Bello 2013). Functional convergence is expected to be higher in harsh environments due to the reduced number of viable ecological strategies and selection for small set of optimal strategies (de Bello et al. 2013; Monge-González et al. 2021). In practice, it is crucial to understand the role of functionally rare species because of their highly original combinations of traits (Mouillot et al. 2013, Violle et al. 2017, Lavergne et al. 2004). Endemic species in particular can be better adapted to their environment of origin in comparison to species with large distribution (Drury 1974). Examples include endemic species in the cuprophilous savannas of central Africa, which have extreme and non-redundant trait values compared to more widespread species (Delhaye 2018), and metal hyperaccumulating plants, often endemic to metalliferous habitats, that tend to have original traits (Gervais-Bergeron et al. 2023) with potentially significant effects on the community.

Intraspecific trait variability (ITV) contributes significantly to overall trait variability in many environments (Albert et al. 2010; Violle et al. 2012; Siefert et al. 2015). ITV is an important driver of community and ecosystem dynamics (Raffard et al. 2019), influences important ecological processes (Ellers et al. 2011; Bolnick et al. 2011) and is essential for assessing eco-evolutionary dynamics, as it plays an important role in the adaptation of individuals and species to environmental pressures (Schoener 2011; Turcotte et al. 2011; Violle et al. 2012). Failing to include intraspecific trait variability in functional diversity indices may lead to erroneous conclusions about community processes (Wong & Carmona 2021), especially in conditions of increased environmental harshness, where its functional importance appears to be higher (Niu et al. 2020). Further, it was suggested that an increase in functional trait coordination could be an adaptive response to environmental stress in plant communities (Dwyer & Laughlin 2017, Delhaye et al. 2020a), but also an intraspecific level response to harsh environmental conditions (Benavides et al. 2021).

Metalliferous soils are scarce but represent good opportunities to study eco-evolutionary processes thanks to the filtering effect of metal toxicity on the vegetation (Harisson & Rajakaruna 2011; Rajakaruna 2018). Serpentine (ultramafic) habitats are the main type of metalliferous environment on the planet, supporting many endemic species (Kazakou et al. 2008; Jakovljević et al., 2022). Serpentine soils are characterized by high concentrations of heavy and trace metals (Ni, Cr, Co, and sometimes Mn and/or Cu), low concentrations of macronutrients (such as N, P and K) and low Ca/Mg molar ratios (Kazakou et al., 2008). Although community assembly processes and the role of intraspecific trait variation have been studied in some metalliferous ecosystems, such as the “copper hills” of the Zambezian region (Delhaye et al. 2016, 2020b), most studies compare trait patterns between serpentine and neighbouring non-serpentine soils (e.g. Fernandez-Going et al. 2012; Adamidis et al. 2014a; Samojedny et al. 2022), highlighting the alternative plant strategies in these contrasting environments. Within metalliferous communities, the relative importance of the processes of community assembly in response to metal toxicity in serpentine soils remains largely unknown. In this study, we tested if serpentine tolerant plant species exhibit intraspecific trait variation in response to metallic stress and we quantified the extent of this variation on six leaf functional traits related to resource acquisition, stress response and competitive ability. Using two complementary approaches, we tested the selective values of these traits and their diversity within communities along a natural metal gradient. Finally, we tested the functional role of the serpentine endemic and Ni-hyperaccumulating species *Odontarrhena lesbiaca* P. Candargy (Brooks et al. 1979; Kazakou et al. 2010) on the communities’ functional composition. Altogether, we expect:

1. Functional convergence in the community toward a stress tolerance syndrome reflecting the strong environmental filter on metal rich soils, and
2. High intraspecific trait variation and covariation and decrease in intraspecific trait diversity reflecting the intensity of the environmental filter represented by the increase in soil metal content.
3. Original trait combination coupled with a unique response of the serpentine endemic and metal-hyperaccumulating species in response to the soil metal content, reflecting its high degree of adaptation to metal toxicity.

## Materials and Methods

### Plant communities and soil

Twenty-six 0.5 m × 0.5 m plots were sampled across four sites (Vatera: n = 7, Ampeliko: n = 5, Olympos: n = 5 and Loutra: n = 9), located in the central and south-eastern part of Lesbos Island (Adamidis et al. 2014b). There is a large variation in soil characteristics within and between sites, originating from variation on the serpentine character (Kazakou et al. 2010), which is reflected by some degree of heterogeneity in the vegetation (Adamidis et al. 2014b).

Plant species richness was determined by visual inspection of each plot, in early June. In each plot, above-ground standing biomass was harvested at the time of peak standing crop. Biomass was oven-dried at 80◦C for 48 h and weighed. Standing biomass was sorted into species. Total above-ground biomass per plot was calculated as the sum of standing biomass and litter and was extrapolated to the scale of 1 m^2^. To characterise the soil chemical composition at the plot level, a soil sample was taken at about 10 cm depth. All soil samples were air-dried and then passed through a 2-mm sieve and stored at 4◦C until analysis. A sub-sample of 4–5 g from each initial soil sample was ground for 70-mesh (<215 µm) sieving and dried at 70◦C and then the total metal concentrations were determined by inductively-coupled plasma emission spectroscopy. Details of the chemical analysis are given in Kazakou et al. (2010).

### Traits measurements

Six foliar functional traits related to plant growth rate, nutrient use, resistance to stress and competitive ability were measured on the youngest fully expanded leaves of the 20 most abundant species across our study sites. All traits were measured with 10 – 30 replicates per species per site where the species were present, using standardized procedures (Pérez-Harguindeguy et al. 2013). Leaf area was measured with a leaf area meter (LI-3050C Transparent Belt Conveyer; LI-COR, Lincoln, NB). Specific leaf area (SLA) was calculated as the ratio of the water-saturated leaf area to the leaf dry mass. Leaf dry matter content (LDMC) was determined as the ratio of leaf dry mass to water-saturated fresh mass and is equivalent to 1 – water content. Leaf thickness (LT) was estimated by the (SLA x LDMC)^−1^ product, or the water-saturated leaf fresh mass to leaf area ratio (Vile et al. 2005). Leaf nitrogen content (LNC) was measured using an elemental analyser (Carlo Erba Instruments, model EA 1108, Milan, Italy). Leaf nitrogen content was not measured on *Bromus commutatus*, *Phleum pratense* and *Trifolium campestre*. Finally, leaf shape is a rarely measured trait that can however be indicative of the plant response strategy to temperature, drought, and light stress because broader leaves (smaller length/width ratio) have a larger layer of immobile air around the stomata (Nicotra et al. 2011, Varsamis et al. 2022). Here, an index of leaf shape was measured on each individual as leaf length/leaf width (unitless). For some sites where the traits of the species could not be measured (8 species-site combinations), the trait values were inferred using a multilevel linear model with no fixed effect and soil Ni as predictor with species identity as random variable (trait ∼ 0 + (1 + Ni | species), using the package lme4 (Bates et al. 2015).

### Statistical analyses

To determine the relationships between environmental variables, we conducted a Principal Component Analysis on the correlation matrix of the log_10_ transformed concentration of all important soil variables (Ni, Cr, Ca, Co, Zn, Mn, Mg/Ca ratio). Since Ni was the environmental variable most correlated with PC1 and is highly correlated with all other toxic elements (Mn, Cr, Co, see Results) it was chosen as the main toxicity gradient in our study. We tested the association between plant biomass and total species richness and the Ni gradient using linear regressions.

Trait values (except SLA and LDMC) were square root transformed before analyses to reduce the effect of extreme trait values on further analyses. Relationships between traits were investigated using a PCA on the average trait value measured in each site.

#### Community trait composition

Recently, Leps & de Bello (2023) emphasized that the adaptive value of traits and their effect on ecosystem processes must be quantified using separate approaches. While the trait adaptive value for each species is better measured using a multilevel model with species as random variables (e.g. Jamil et al. 2013, ter Braak 2019), the classical approach of regressing the community weighted mean trait values along environmental gradients (CWMr) is informative of the community effect on ecosystem processes. Here, we used both these approaches as complementary. We used the MLM3 method (ter Braak 2019) to investigate the selective value of each trait, as it provides a better control of type I error and a higher power compared to other previously proposed multilevel model approaches. We used square root transformed measures of abundance rounded to the closest upper integer, to avoid transforming small abundance values into false zeros. We fit a multilevel model with the interaction between trait and soil Ni content as a fixed effect and two random effects, 1) the effect of trait per site and 2) the effect of soil Ni content per species, using a Poisson distribution of the response variable (ter Braak, 2019). The models were computed in a Bayesian framework in STAN using the *brms* package (Bürkner 2017) with 4000 iterations and vague priors.

To measure the dominant trends in trait values and the potential effect on ecosystem functioning, we calculated the community weighted mean (CWM, Lavorel & Garnier 2002) of each trait in each plot. We also investigated the relative importance of different community assembly processes using the functional dispersion index (FDis, Laliberte & Legendre 2010). FDis measures the average relative distance of each species to the centroid of trait values in a multivariate space, weighted by species relative abundance. Since multivariate indices of functional diversity could hide specific patterns linked to some axes of functional variation (Butterfield & Suding 2013), we calculated both multivariate and the univariate sesFDis index on each trait separately. Because functional diversity metrics are sensitive to the number of species in the community, we used a null model approach to calculate the standardised effect size of the FDis index (sesFDis) as described in Delhaye et al. (2020b). For each plot, we permutated the abundance of each species (n = 500) and recalculated FDis each time to produce a null distribution of the index. We extracted the mean and standard deviation of this distribution and calculated the 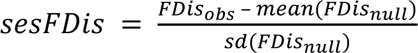. Finally, to test the effect of the serpentine endemic *O. lesbiaca* on the functional characteristics of the communities, we repeated these analyses after excluding that species from the species pool. The variation of CWM, and sesFDis along the gradient of soil Ni content was tested using a linear model.

#### Intraspecific trait variation

To assess the relative contribution of intra and interspecific trait variation to the total community level variation, we used a variance partitioning method based on a random intercepts and slopes of a multilevel linear mixed model. We built a model of the traits distribution with a fixed effect of the soil Ni gradient and a random effect with varying intercept and slope for each species using the *brms* package. We then decomposed the variance of the model into three components: 1) interspecific, explained by the intercept of the random effect, 2) intraspecific explained by the varying slope of the random effect and, 3) the residual intraspecific variation. We used the same model to investigate the slope of the relationship between trait and soil Ni gradient for each species separately. To quantify the effect of intraspecific variation on community processes and the relative importance of intra and interspecific variation and their covariance to the measure of CWM, we used the CWM decomposition method proposed by Lepš et al. (2011). Finally, we tested for a change in intraspecific trait hypervolume, functional dispersion (FDis) and coordination for each site along the Ni gradient. To quantify functional coordination, we used the variance of the eigenvalues of a correlation matrix between traits of a species in a specific environment (Cheverud et al. 1989, Boucher et al. 2013). We regressed these three indices along the soil Ni gradient using a multilevel linear model with gamma distribution and a random intercept and slope for each species, using the glmer function.

All analyses were conducted in R (R Core Team 2022) using R Studio (R Studio Team 2020) and tidyverse (Wickham et al. 2019) for data manipulation and figure creation.

## Results

### Soil and plant community

Soil Ni content ranges from 210 to 3740 mg kg^−1^ and soil Cr content ranges from 143 to 7065 mg kg^−1^. There is a strong correlation between both metals in the soil (r = 0.82). Other potentially toxic metals in the soil show a lesser extent of variation: Mn (668 – 2303 mg kg^−1^), Co (2 – 135 mg kg^−1^), Zn (53 – 148 mg kg^−1^) and are highly correlated to the soil Ni content (r = 0.93; 0.91 and 0.85 respectively, Figure 1). The Mg/Ca ratio varies from 2.6 to 12.3 and is only moderately positively correlated to soil Ni content (r = 0.56).

**Figure 1.**
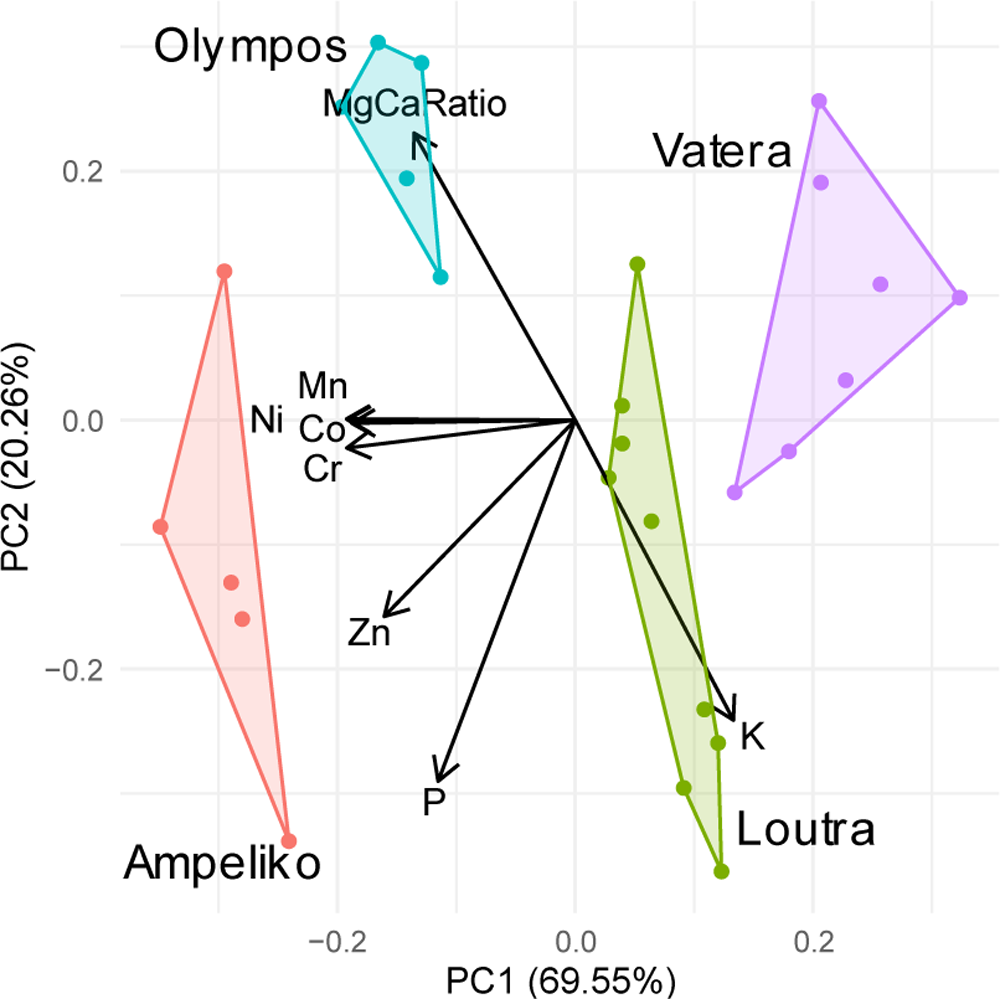
Biplot of the principal component analysis on correlation matrix of the standardised soil variables.

The first two principal components explain 90% of the variance. PC 1 is mostly correlated to Ni, Cr, Mn, and Co, while PC2 is mainly correlated to P, K, and the ratio Mg/Ca (Figure 1), suggesting that metal toxicity and nutrient content are independent in the studied plots. The four sites are clearly separated on the biplot with Ampeliko showing the strongest serpentinic characteristics, followed by Olympos, Loutra and finally Vatera.

Species richness in each plot varies from 15 to 38 with a biomass of 102 to 374 g/m^2^. There is no significant change in species total richness, or biomass along the Ni gradient.

### Community level variation

There is good evidence that a larger leaf thickness increases species abundance with increasing soil Ni content (p _β > 0_ = 0.95). However, for all other traits there is no strong indication that their value influences the abundance response along the Ni gradient (Fig. 2)

**Fig 2.**
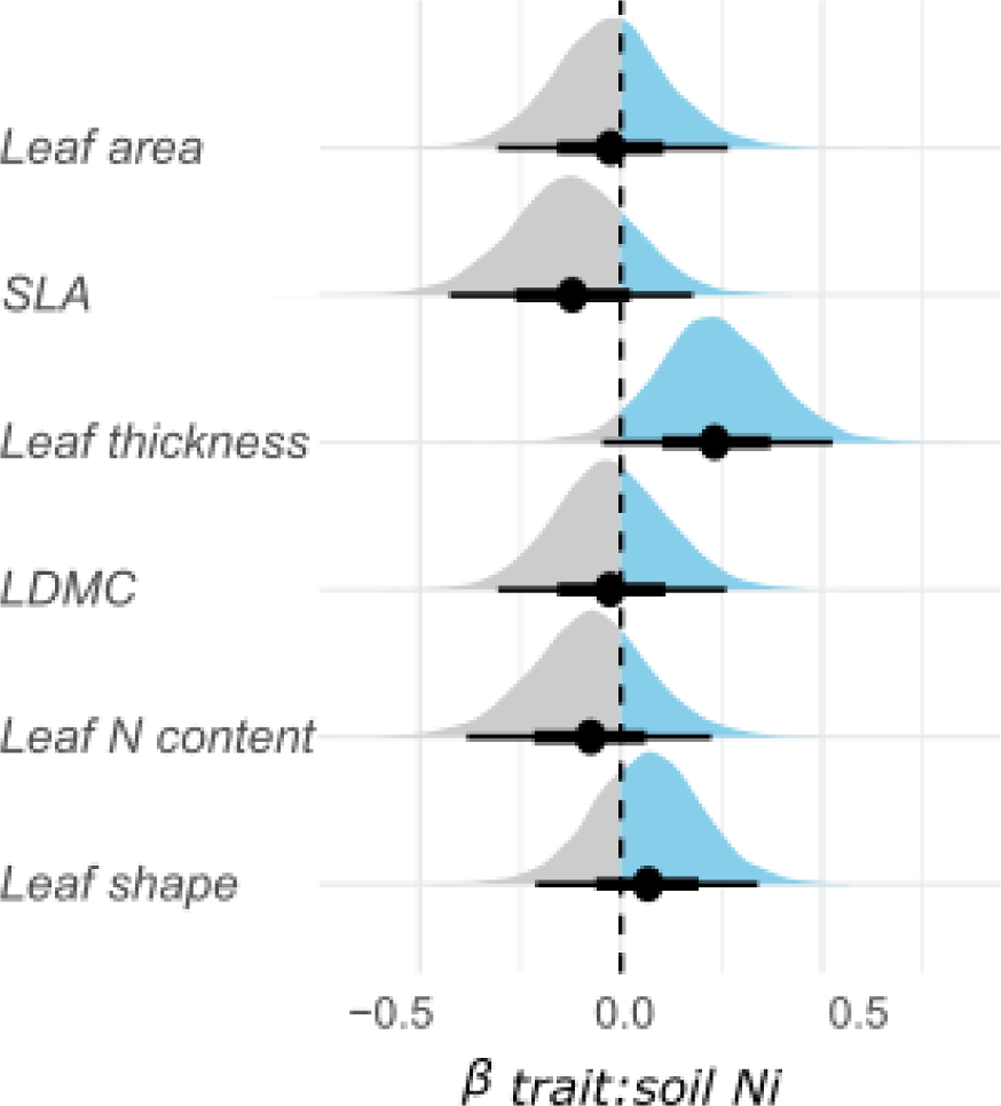
Density plot of the posterior distribution of the interaction betweem soil Ni and trait values in our communities. Positive values indicate that the abundance of species with a higher trait values increases when increasing soil Ni content.

The community weighted means including intraspecific trait variation show contrasting patterns depending on the trait and the presence of *O. lesbiaca* (Fig. 3). The complete communities exhibit a decrease in SLA, and an increase in leaf thickness with increasing soil Ni content. Leaf dry matter and shape do not show any clear pattern. When removing *O. lesbiaca* from the community, these two relationships disappear, while leaf area and specific leaf area increases significantly with increasing soil Ni content.

**Figure 3.**
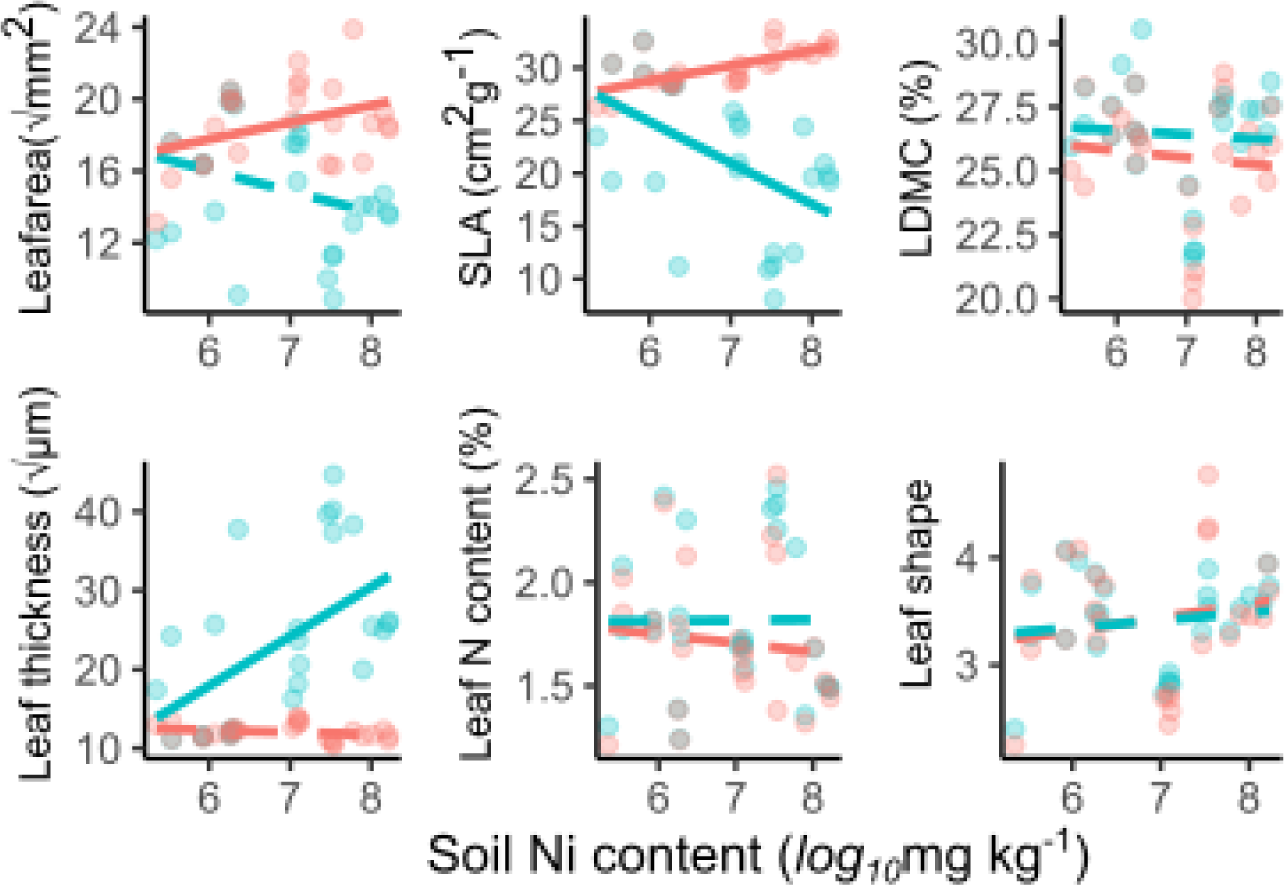
Variation of the community weighted mean value for each trait (including intraspecific trait variation) along the Ni gradient. The blue line and points represent the whole community, the red line represents the community without *O. lesbiaca* (see methods). Plain lines are significant and dashed lines are non-significant regression lines (α < 0.05).

Regarding functional dispersion, the multivariate sesFDis does not show any significant trend (Fig. S2). Univariate functional dispersion shows trait specific patterns (Fig. 4). In the complete community dataset, the diversity of SLA and leaf thickness are over-dispersed while leaf shape tends to be clustered at the high Ni end of the gradient. Leaf dry matter and nitrogen contents do not show any clear pattern. When removing *O. lesbiaca* from the community, all significant relationships disappear.

**Figure 4.**
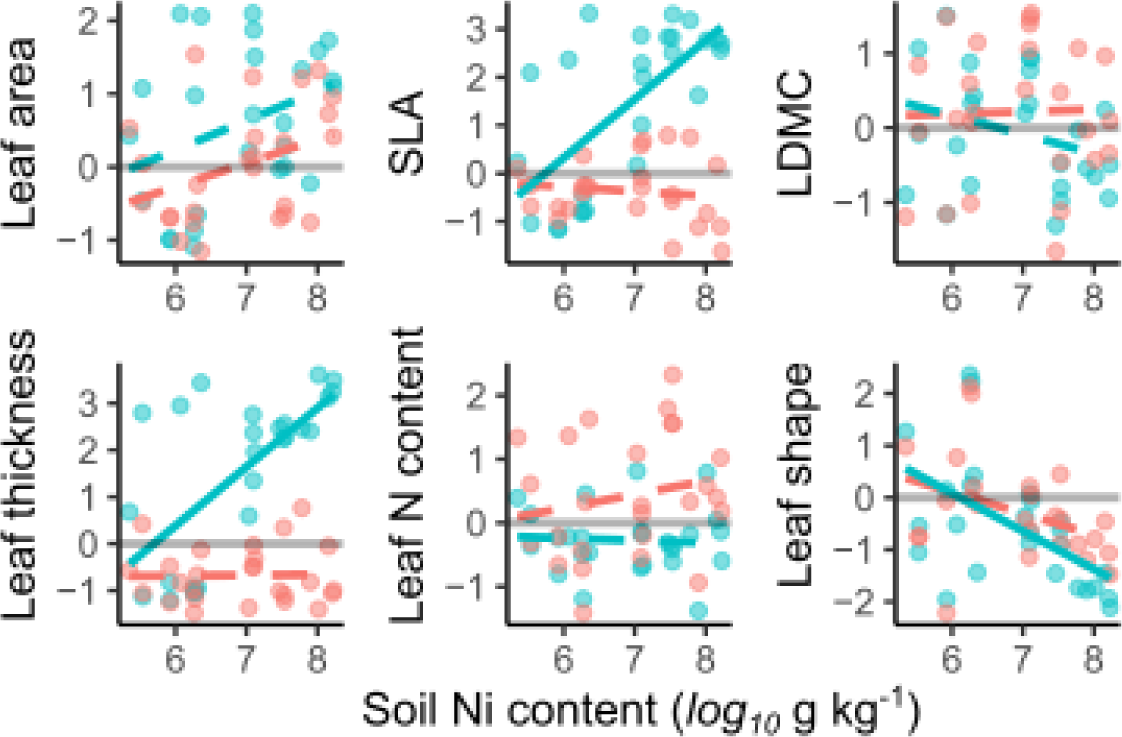
Standardised effect size of the univariate functional dispersion (FDis) for each of the six studied traits (including intraspecific variation) along the soil Ni content gradient. The blue line and points represent the whole community, the red line represents the community without *O. lesbiaca* (see methods). Plain lines are significant and dashed lines are non-significant regression lines (α < 0.05). The grey horizontal line represents the zero intercept.

### Intraspecific variation

For all traits, interspecific differences are much larger than the intraspecific variation explained by the Ni gradient (Fig. 5). The interspecific differences in traits account for between 38% of the variance for SLA and leaf N content and 92 % of the total variance for leaf thickness. For leaf N content and SLA, most of the variation is explained by the residual intraspecific variation. For all traits, the ITV explained by the Ni gradient is smaller than 2%.

**Figure 5.**
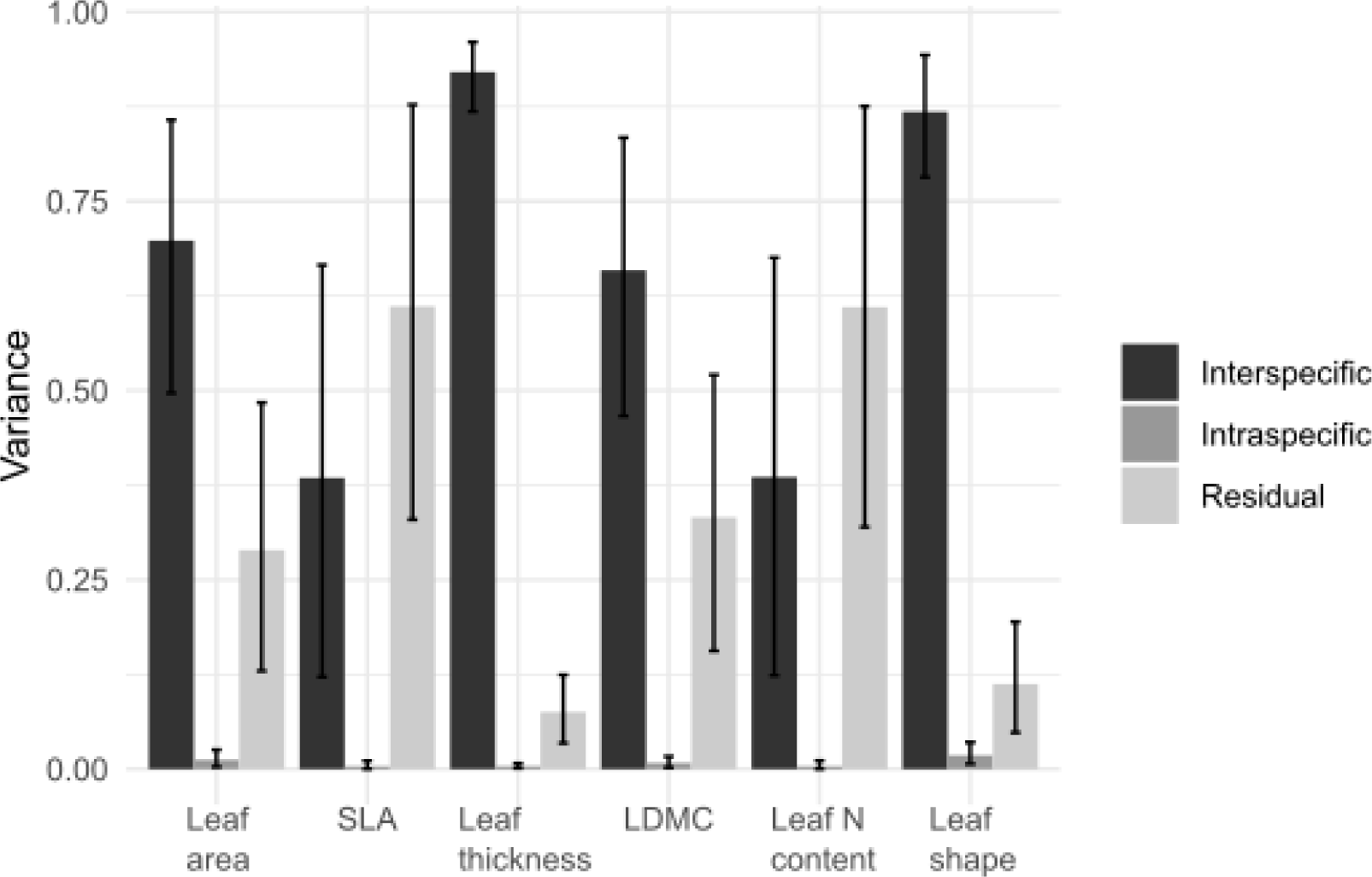
Partitioning of the variance between interspecific differences, intraspecific variation explained by the soil Ni gradient and the residual unexplained intraspecific variation. Each value is the median value of the posterior distribution, and the incertitude represents the 95% credibility interval for each parameter.

Using the method based on the variance decomposition of CWM calculated including or excluding ITV, the interspecific differences in trait are much larger than intraspecific except for LDMC, which only explains a minuscule part of the variation overall (Fig. S3). Interestingly, for all traits, the covariation between the intra and interspecific fractions is negative, suggesting that species responses are in the opposite direction compared to the community trends.

No general patterns emerged in the estimate of intraspecific trait variations. Most species did not show significant trait variation between low and high nickel sites (Table 1). Along the soil Ni gradient, two species (*A. barbata* and *C. commutata*) show a clear increase in leaf area, and three species (*O. lesbiaca*, *A. arvensis* and *T. arvense*) show a decrease while one species (*F. eriocephala*) shows an increase in leaf thickness. In addition, *D. glomerata* shows a decrease and *L. rigidum* shows an increase in leaf dry matter content while *L. rigidum* shows an increase and *A. barbata* shows a decrease in leaf length/width ratio. No species shows a clear trend in SLA and N content values along the soil Ni gradient.

**Table 1.** Slope and associate 95% credibility intervals for the traits of each species along the soil Ni gradient, obtained using a multilevel model of trait variation with species as random variable. Slopes values where the CI does not intercept zero are shaded. Nomenclature for plant species follows Flora Europea as has been specified in Adamidis et al. 2014b.

| Species | Leaf area | SLA | Leaf thickness | LDMC | Leaf N content | Leaf shape |
| --- | --- | --- | --- | --- | --- | --- |
| *Aegilops biuncialis* | -0.43 (-1.59 : 0.69) | 0.79 (-0.51 : 3.65) | -0.49 (-1.37 : 0.31) | 3.43 (-8.83 : 15.93) | 0.79 (-0.5 : 3.62) | 0.03 (-0.19 : 0.24) |
| *Odontarrhena lesbiaca* | 0.4 (-0.69 : 1.54) | -0.51 (-2.88 : 1.57) | -1.23 (-2.07 : -0.38) | -4.53 (-17.14 : 7.4) | -0.48 (-2.86 : 1.62) | 0.09 (-0.11 : 0.3) |
| *Anagallis arvensis* | 0.24 (-0.91 : 1.52) | -0.13 (-2.02 : 1.84) | -1.61 (-2.59 : -0.72) | 7.34 (-6.66 : 22.68) | -0.11 (-2.05 : 1.85) | -0.13 (-0.36 : 0.09) |
| *Avena barbata* | 2.67 (1.25 : 4.16) | -0.09 (-2.09 : 1.79) | 0.32 (-0.57 : 1.21) | -1.67 (-15.63 : 11.94) | -0.09 (-2.06 : 1.65) | -0.32 (-0.57 : -0.09) |
| *Crepis commutata* | 1.36 (0.11 : 2.79) | 0.21 (-1.89 : 2.38) | 0.33 (-0.58 : 1.28) | -6.83 (-22.71 : 7.97) | 0.19 (-1.94 : 2.33) | 0 (-0.24 : 0.25) |
| *Cynosurus echinatus* | 0.18 (-1.28 : 1.74) | 0.06 (-2.23 : 2.14) | 0.58 (-0.56 : 1.83) | -3.19 (-21.66 : 12.3) | 0.05 (-2.38 : 2.12) | 0.15 (-0.17 : 0.5) |
| *Dactylis glomerata* | -1 (-2.25 : 0.09) | 0.16 (-1.39 : 2.2) | 0.49 (-0.33 : 1.29) | -13.28 (-28.37 : -0.03) | 0.15 (-1.41 : 2.13) | 0.1 (-0.12 : 0.31) |
| *Filago eriocephala* | -0.35 (-1.53 : 0.79) | -0.68 (-3.16 : 0.99) | 0.87 (0.04 : 1.76) | -9.03 (-22.87 : 3.37) | -0.67 (-3.11 : 0.93) | -0.04 (-0.27 : 0.19) |
| *Hordeum bulbosum* | 0.06 (-1.02 : 1.11) | 1.04 (-0.24 : 3.88) | -0.3 (-1.08 : 0.45) | 3.76 (-8.29 : 16.22) | 1.05 (-0.31 : 3.95) | 0.03 (-0.18 : 0.24) |
| *Lagoecia cuminoides* | -1.02 (-2.27 : 0.07) | -0.03 (-2.03 : 1.61) | 0.28 (-0.53 : 1.11) | -5.09 (-18.28 : 7.86) | -0.05 (-2.03 : 1.51) | -0.01 (-0.23 : 0.21) |
| *Lolium rigidum* | 0.27 (-0.81 : 1.34) | -0.44 (-2.9 : 0.83) | 0.18 (-0.6 : 0.94) | 22.92 (9.1 : 38.26) | -0.46 (-3.05 : 0.77) | 0.95 (0.73 : 1.18) |
| *Phleum pratense* | -0.21 (-1.36 : 0.88) | 0.03 (-2.08 : 1.72) | 0.15 (-0.64 : 0.95) | 3.43 (-8.82 : 15.85) | 0.01 (-2.09 : 1.71) | -0.18 (-0.4 : 0.04) |
| *Plantago lagopus* | -0.57 (-1.78 : 0.52) | -0.02 (-1.87 : 1.7) | 0.47 (-0.32 : 1.3) | 4.48 (-8.42 : 18.41) | -0.03 (-1.9 : 1.66) | -0.14 (-0.36 : 0.07) |
| *Sanguisorba minor* | -0.13 (-1.25 : 0.95) | 0.05 (-1.8 : 1.81) | 0.34 (-0.43 : 1.16) | -5.41 (-18.66 : 6.85) | 0.04 (-1.79 : 1.73) | -0.05 (-0.26 : 0.17) |
| *Torilis nodosa* | -0.44 (-1.74 : 0.78) | -0.3 (-3.1 : 1.11) | 0.51 (-0.38 : 1.48) | 0.8 (-13.94 : 15.79) | -0.3 (-3.14 : 1.11) | -0.04 (-0.29 : 0.21) |
| *Trachynia distachya* | -0.35 (-1.49 : 0.74) | -0.46 (-2.99 : 0.79) | 0.03 (-0.78 : 0.83) | 6.15 (-5.91 : 19.3) | -0.44 (-3.15 : 0.83) | -0.02 (-0.24 : 0.2) |
| *Trifolium angustifolium* | -0.13 (-1.36 : 1.1) | 0.56 (-0.82 : 3.73) | -1.24 (-2.25 : -0.32) | 4.01 (-9.45 : 18.37) | 0.54 (-0.8 : 3.54) | -0.14 (-0.38 : 0.09) |
| *Trifolium arvense* | -0.49 (-2.58 : 1.49) | 0 (-2.35 : 2.43) | 0.07 (-1.57 : 1.75) | -0.06 (-21.52 : 20.36) | -0.01 (-2.59 : 2.28) | -0.18 (-0.76 : 0.38) |

There was no clear pattern of change in intraspecific trait coordination, richness, or dispersion along the soil Ni gradient.

## Discussion

### Community-level trait variation

At the community level, leaf thickness is the main trait driving species abundance along the Ni gradient. These interesting results could be linked to the fact that low water availability on serpentinic soils most likely exerts a strong selective force on the vegetation (Brady & Kurckeberg 2005). This absence of selective value detected for other traits could also be due to the fact that all our sites are serpentinic, and only vary in degree of soil metal toxicity. Therefore, plant species growing in these environments are most likely adapted to metal toxicity and a stronger filtering effect of trait values could be observed when comparing these sites with non-serpentinic communities. However, we cannot rule out that the specific Ni gradient studied here, might not have any direct effect on the measured traits. Indeed, a common hypothesis is that there is a trade-off between metallic stress tolerance and resource capture, growth, and competitive ability (the “cost of tolerance”, Kruckeberg, 1951; Wilson, 1988). However, our results do not support this hypothesis.

The community showed a trend towards lower SLA and thicker leaves (Fig. 3) with increasing soi Ni. This is coherent with the hypothesis that traits of the “leaf economics spectrum” converge towards a stress tolerant strategy in stressful environments (Wright et al. 2004). However, most of the community trait variation is driven by the traits of *O. lesbiaca*. When *O. lesbiaca* is removed from the community, the remaining species do not show an increase in SLA and leaf are values in response to soil nickel. The increase in SLA is surprising, considering that a stress tolerant strategy would imply lower SLA (Wright et al. 2004), but this is probably due to the dominance of annual herbaceous species (Kazakou et al. 2010; Adamidis et al. 2014b) in the floristic composition of these Mediterranean communities. Delhaye et al. (2020a) hypothesized that the rapid growth of annuals within a short life cycle on soils with high metal contents promotes metal accumulation, while the slow growth and longer life cycles of perennials lead to long-term strategies of metal exclusion from plant tissues. This result is coherent with what is observed on copper hills (Delhaye et al. 2016) and with *Noccaea* (*Thlaspi*) *caerulescens* across Europe (Dechamps et al. 2007). These changes in dominant trait values along the gradient could influence ecosystem processes such as litter decomposition (Kazakou et al., 2006) and interactions with soil fauna (Lam et al. 2024).

The absence of clustering for the functional diversity index along the whole gradient, likely reflects the fact that the species present have highly conserved trait values that do not respond to metal toxicity. To study the effect of environmental filters on trait combinations, the study of interspecific trait coordination (Delhaye et al. 2020a) could provide a complementary approach. However, this so far relies on differences in community composition and here we selected specifically the species present along most of the gradient, making it impossible to detect any change in interspecific coordination.

### Low intraspecific trait variation in response to soil metal content

We expected to find strong intraspecific variation for some leaf traits coherent with a stress tolerance and resource conservation strategy, as observed recently for the facultative metallophyte *Silene paradoxa* L. (Lazzaro et al. 2021). Our results showed that intraspecific trait variation in response to the toxicity gradient was low compared to interspecific differences (Fig.2). ITV was lower than global estimates measured across communities globally (Siefert et al. 2015), but similar to what has been observed in other metalliferous environments such as the copper hills of the Zambezian region in Africa (Delhaye et al. 2016). Intraspecific variability has been shown to be trait-specific (e.g., Auger & Shipley 2013; Delhaye et al. 2016, 2018), and it is generally low in leaf traits such as SLA (Kazakou et al. 2014) or shape that are more phylogenetically conserved (Siefert et al. 2015).

With increasing metal stress, some species tend to follow more conservative resource use strategies based on the predictions of the leaf economics spectrum (Wright et al. 2004; Kazakou et al. 2008; Adamidis et al. 2014a). For example, in nickel-rich soils the leaves of *O. lesbiaca, A. arvensis and T. angustifolium are* significantly thicker and *L. rigidum* has higher leaf dry matter content and length/width ratio. These changes could indicate a response to drought stress on soil with high Ni content. However, these responses are not generalised. Some species show an increase in leaf area which could indicate better growth on high Ni content soils. This could be due to a lower pathogen pressure on metal rich soils (Boyd et al. 1994; Boyd 2007), which could favour plant growth. Other physiological traits, such as proline content, might help disentangle the effects of drought and metal toxicity. Proline content plays a crucial role in osmoregulation (Verbruggen & Hermans 2008) and is higher in plants grown on serpentine than non-serpentine soils (Karatassiou et al. 2021). This could represent an interesting functional trait for investigating the response of plant species to metallic trace elements. Even though N shows a large intraspecific variation, other crucial nutrients like Ca, Fe and P content, could also show interesting patterns of variation in response to soil toxicity (Kazakou et al. 2008, 2010; Karatassiou et al. 2021), and should be investigated further. Finally, for species showing a significant intraspecific response in some traits, it would be interesting to test if this difference in traits is due to phenotypic plasticity or to local adaptation (Dechamps et al. 2007; Jimenez-Ambriz et al. 2007; Rajakaruna 2018), even if the unfavourable environmental conditions reduce the possibility of expression of phenotypic plasticity patterns (Huang et al. 2015). Trait variability can, however, be shaped by abiotic factors other than soil elemental concentrations such as soil water retention or biotic interactions (reduced competition or pathogens) on metal rich soils, and therefore, experimental approaches should be used. We did not find any change in intraspecific trait coordination and diversity along the Ni gradient. A possible explanation for this result is that all the traits used here are related to resource capture and conservation, but none of them has a causal link to metal tolerance. In the future, we suggest combining different functional axes, notably foliar traits related to resource capture, root traits and chemical traits related to metal tolerance strategy (Bert et al. 2002).

### The functional originality of O. lesbiaca

It was suggested that strict metallophytes (edapho-endemic) could possess original and more extreme trait values than widespread facultative metallophytes (Delhaye 2018). Compared to the other species in the community, *O. lesbiaca* has original trait values and combinations, suggesting that endemic species can be functionally unique compared to widespread species in the Mediterranean region (see Lavergne et al. 2003). This is likely the result of evolution towards traits of resistance to metal linked to hyperaccumulation (Kazakou et al. 2008). *O. lesbiaca* is the only hyperaccumulator species in our communities, with differences in Ni hyperaccumulation among its populations being consistent with nickel availability in the soil (Kazakou et al. 2010; Adamidis et al. 2014c). In a rhizotron experiment, individuals of *O. lesbiaca* from Ni rich soils (Ampeliko) proved to be more tolerant in terms of root growth relative to those from sites with lower nickel content (Loutra), as Ni-induced nitric oxide accumulation was observed in their root tips along with a decrease in the degree of protein tyrosine nitration (Feigl et al. 2020). The difference in traits of the *O. lesbiaca* could also be linked to high resistance to low nutrient and water availability (Kazakou et al. 2008; Harrison et al. 2017), as species strictly restricted to serpentine habitats are found to have higher drought tolerance than more generalist Mediterranean species (Hidalgo-Triana et al. 2023). The link between metal hyperaccumulation and drought resistance has been hypothesized (Severne, 1974): excess nickel in cells can act as an osmoticum, reducing the water potential in plants (Baker & Walker, 1989), and allowing photosynthesis to continue during drought conditions (Fitter & Hay, 1981).

Finally, *O. lesbiaca* is a subshrub with a woody base (life form chamaephyte), while most of the other studied species are therophytes or hemicryptophytes (FloraVeg.EU, 2023), which have shorter life spans and faster growth rates and thus different trait values (Garnier et al. 1997; Samojedny et al. 2022). These extreme traits make it difficult to infer community assembly processes from the traits of the whole community, and this could be the case in many stressful environments where there could coexist different combinations of trait values in response to each specific stressful condition.

## Conclusions

Intraspecific variation in leaf traits responds little to a large soil Ni gradient relative to the differences in trait values between species. Endemic plant species that hyperaccumulate metals may have original trait values compared to species with a wider distribution, which reinforces the need for their conservation. Future studies should focus on the relative contribution of species turnover and intraspecific variation to trait shifts across the continuum from resource acquisition to stress tolerance strategies along large gradients of soil toxicity to highlight the ecological and evolutionary mechanisms leading to such original plant communities.

## Acknowledgements

GCA received funding from the programme MEDICUS of the University of Patras. GD was supported by the Philippe Wiener–Maurice Anspach Foundation.

## Author contributions

GD, GCA and PGD developed the ideas, GCA and PGD collected the data, GD analysed the data. All authors contributed to the redaction and revision of the manuscript.

## Data availability

The R code and data used for the analyses are available at github.com/guildelhaye/serpentine_community.

