## Supplementary material for "Interspecific trait differences drive plant community responses on serpentine soils"

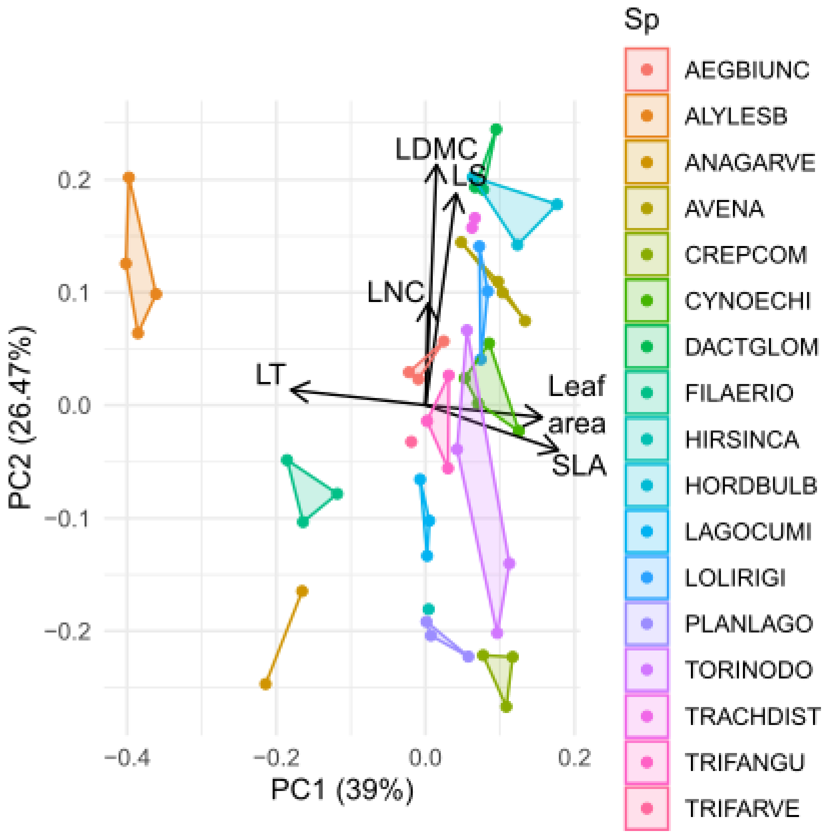


Figure S1. Principal component analysis of trait values of the 17 studied species in each site where traits were measured. Most species show a limited trait variation between sites.

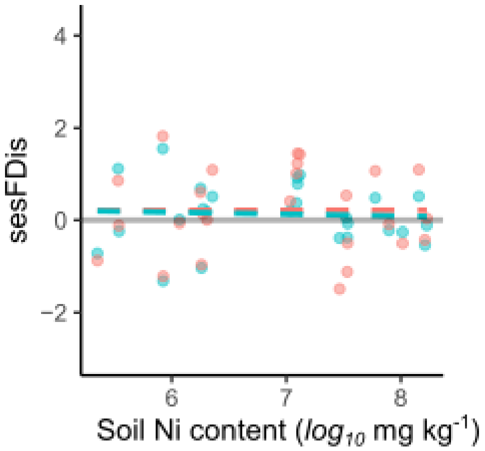


Figure S2. Standardised effect size of the multivariate index of functional dispersion (FDis) along the soil Ni content gradient. The blue line and points represent the whole community, the red line represents the community without *O. lesbiaca* (see methods). Both regressions are non-significant. The grey horizontal line represents the zero intercept. Points above the intercept are communities showing functional overdispersion, points under show functional clustering.

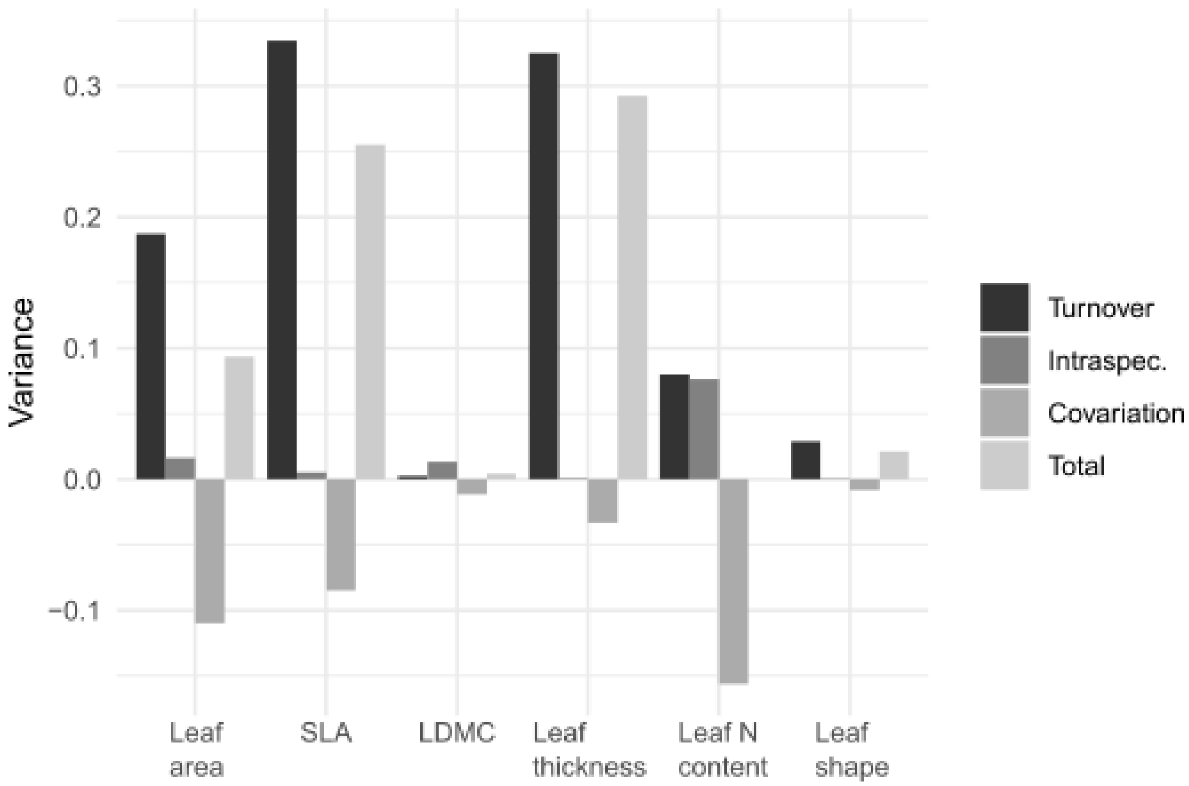


Figure S3. Variance decomposition of a linear model of the community weighted mean of each trait value along the soil Ni gradient. The total sum of square is decomposed into interspecific, intraspecific and covariation components using the method proposed in Lepš et al. (2011).
